## Supplementary figures and images for "Fecal microbiota transplantation derived from Alzheimer’s disease mice worsens brain trauma outcomes in wild-type controls"

### Graphic Abstract

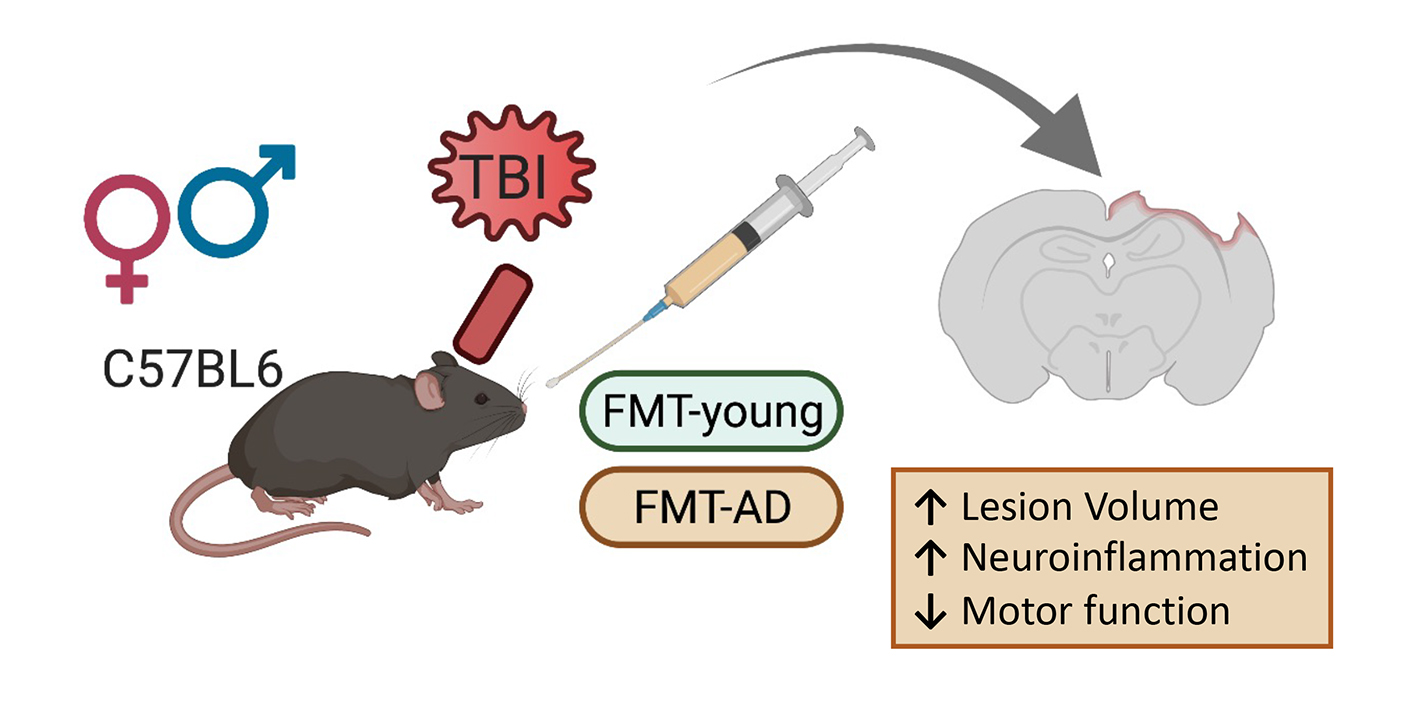
